# Donor-derived vasculature is required to support neocortical cell grafts after stroke

**DOI:** 10.1101/2021.02.27.433204

**Authors:** Joanna Krzyspiak, Jingqi Yan, Hiyaa Ghosh, Basia Galinski, Pablo J. Lituma, Karina Alvina, Samantha Kee, Marta Grońska-Pęski, Yi De Tai, Kelsey McDermott, R. Suzanne Zukin, Daniel A. Weiser, Pablo E. Castillo, Kamran Khodakhah, Jean M. Hébert

**Affiliations:** Department of Neuroscience, Albert Einstein College of Medicine, Bronx, USA; Stem Cell Institute, Albert Einstein College of Medicine, Bronx, USA; Department of Genetics, Albert Einstein College of Medicine, Bronx, USA; Department of Pediatrics, Albert Einstein College of Medicine, Bronx, USA; University of Rochester, Rochester, USA; Department of Psychiatry and Behavioral Sciences, Albert Einstein College of Medicine, Bronx, USA

**Keywords:** neocortex, transplantation, ischemia, vascular endothelial cells, blood vessels

## Abstract

Neural precursor cells (NPCs) transplanted into the adult neocortex generate neurons that synaptically integrate with host neurons, supporting the possibility of achieving functional tissue repair. However, poor survival of transplanted NPCs greatly limits efficient engraftment. Here, we test the hypothesis that combining blood vessel-forming vascular cells with neuronal precursors improves engraftment. By transplanting mixed embryonic neocortical cells into adult mice with neocortical strokes, we show that transplant-derived neurons synapse with appropriate targets while donor vascular cells form vessels that fuse with the host vasculature to perfuse blood within the graft. Although all grafts became vascularized, larger grafts had greater contributions of donor-derived vessels that increased as a function of their distance from the host-graft border. Moreover, excluding vascular cells from the donor cell population strictly limited graft size. Thus, inclusion of vessel-forming vascular cells with NPCs is required for more efficient engraftment and ultimately for tissue repair.

## Introduction

After cell loss due to injury or disease such as stroke, neurons in the adult mammalian neocortex do not regenerate, often resulting in devastating functional deficits. Various stem cell types, including neural, mesenchymal, hematopoietic, and endothelial cells, have been introduced into the brain parenchyma as potential forms of therapy in both pre-clinical and clinical studies (Rikhtegar et al., 2019). The vast majority of these studies have shown that the mechanism by which the transplants improve recipient function, if at all, is through a bystander effect –whereby transplanted cells instigate neuroprotection of surrounding host tissue rather than provide a new substrate for neural circuit repair (Zhang et al., 2020). However, in mouse models, transplanted neural precursors can generate neurons that project to appropriate brain targets and electrophysiologically connect with host neurons (Hébert and Vijg, 2018). Consequently, transplant-derived neurons can respond to sensory input such as light and touch, or affect motor activity, supporting the possibility of functional repair (Falkner et al., 2016; Linaro et al., 2019; Michelsen et al., 2015; Palma-Tortosa et al., 2020; Tornero et al., 2017).

Nevertheless, to achieve clinically relevant tissue repair paradigms, several advances in transplantation technology are required. First and foremost, NPC survival after transplantation will need to be greatly improved, especially given the relatively large infarct sizes in humans and the anticipated need for correspondingly large grafts. Transplants of NPCs generally have a low survival rate when administered as a purified cell suspension (e.g. Li et al., 2014; Nakagomi et al., 2009; Hicks et al., 2009), which is the current pre-clinical and clinical norm. This poor survival calls on the field to determine how we can improve the survival of transplanted NPCs to produce large functional grafts. A potentially important process for successful engraftment is vascularization. For instance, during fetal development, vascularization must meet the physiological demands of each growing tissue, including the brain (Vasudevan et al., 2008; Paredes et al., 2018). Similarly, vascularization of neural cell transplants might also need to occur to promote optimal neuron survival, differentiation, and function (Hill and Cave, 2015).

Transplants of human NPCs or cerebral organoids into mouse neocortices exhibit significant cell death until they eventually become vascularized by the host, consistent with the importance of vascularization for graft cell survival (Basuodan et al., 2018; Mansour et al., 2018). However, attempts to increase the survival of transplanted NPCs by including vascular endothelial cells in the donor cell population have thus far been inconclusive or have not shown substantial vessel formation (Cai et al., 2015; Li et al., 2014; Hsueh et al., 2015; Nakagomi et al., 2009; Pham et al., 2018). Instead of NPCs, the transplantation of fetal brain cells or tissue, which include their own vascular cells, yielded unclear results regarding the origin of the vessel-forming cells within the grafts and did not provide evidence for the potential benefits, if any, of donor-derived vessels (Broadwell et al., 1990; Dusart et al., 1989; Krum et al., 1988; Geny et al., 1994; Pennell et al., 1997; Péron et al., 2017). Hence, because no study to date has shown robust development of blood vessels from dissociated donor cells, the effect of including vessel-forming donor cells within transplants remains an open question.

Here, cell transplantations were performed in adult mice with ischemic stroke as an injury model. We chose distal medial cerebral artery occlusion (dMCAO) to confine the stroke damage to the neocortex. We found that transplants of dissociated cells from E12.5 mouse neocortices generate large grafts (up to 15 mm^3^) containing upper and lower cortical neuron identities, often arranged in pseudo-layers, that can project to appropriate host targets and become synaptically integrated. Grafts also formed donor-derived vessels that anastomosed (fused) with host vessels to perfuse blood. Graft size increased with the proportion of total vasculature that came from the donor, with the contribution of donor-derived vessels within the graft increasing as a function of distance from the graft-host border. Interestingly, graft size was also greater when cells were transplanted at stroke sites than when transplanted in uninjured neocortex, suggesting that stroke sites are especially conducive to donor-derived vascularization of the graft. Finally, removing vascular cells from the donor cell population imposed a strict size limit (3 mm^3^) on the grafts. Our findings, therefore, support the hypothesis that combining neural stem cells with vascular cells is necessary to generate large grafts, particularly at ischemic stroke sites.

## Results

### Grafts derived from dissociated fetal neocortices contain synaptically integrated neurons

As a source of dissociated donor cells for transplantation, we chose E12.5 mouse neocortex because it is comprised of neural, vascular, and microglial cells (Loo et al., 2019), providing a source of precursors for potential neuronal and vascular integration with the host. Note that at this stage of development, most cells immunopositive for SCA1, a marker for vascular progenitors, are present in the developing meninges, namely the inner pia, (Supplementary Figure 1), which were included in the preparation of the dissociated donor cell population. For each experiment, the single cell nature of the donor cells was confirmed under lightfield microscopy. Initially, to assess the overall health of grafts, we examined their neurons. In a first set of experiments, donor embryos carried a conditional Tau-promoter-driven membrane-bound-GFP reporter and a *Foxg1^Cre^* allele that marks donor-derived neurons and enables tracing of their projections. Cells were transplanted into stroke sites caused by dMCAO. A 7-day delay between induction of stroke and transplantation was introduced in the timeline (Figure 1A), because this delay after cortical injury favors overall graft vascularization, health, and differentiation (Péron et al., 2017), and because this may correspond to a clinically relevant early time point for transplant treatments after stroke. In 67% of grafts (10 out of 15 grafts), layer-like structures were observed by 5 days post-transplant (dpt) (Figure 1B). Cells labeled with CTIP2, a marker for deep layer cortical neurons, were located further from the center of the grafts than proliferating Ki67+ progenitors and closer to the center than cells labeled for SATB2, a marker for superficial cortical neurons, mimicking the arrangement of a developing cortex. At 14 dpt, CTIP2+ and SATB2+ cells remained present in a pseudolayered cytoarchitecture (Figure 1C).

**Figure 1.**
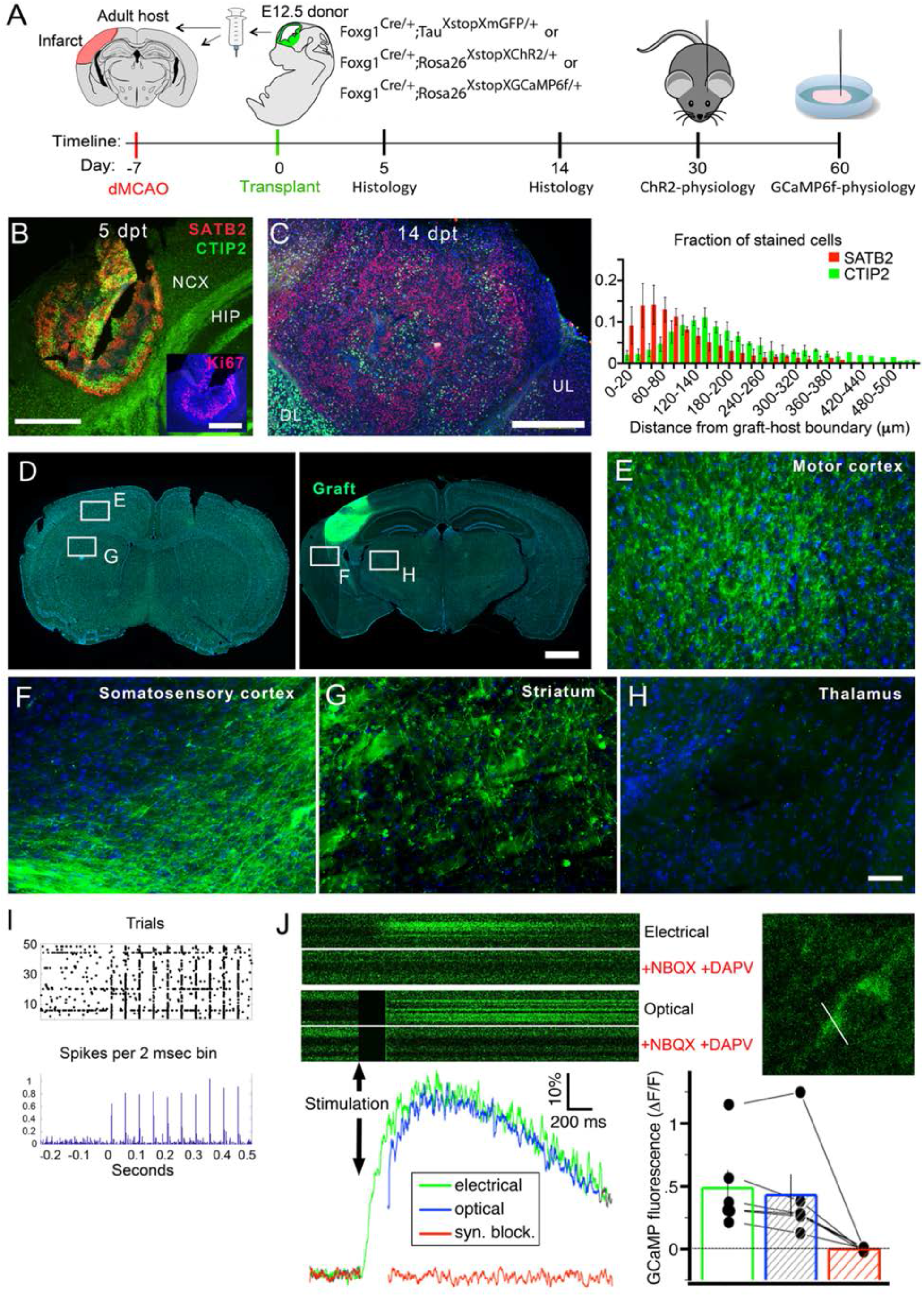
Transplanted E12.5 cortical cells generated upper and lower layer neurons that project and synapse with appropriate targets. A. Experimental timeline. B. Transplanted cells at 5 dpt stained with upper layer marker SATB2, lower layer marker CTIP2, and in the inset at half the magnification proliferation marker Ki67. Scale bars: 400 um. C. Transplanted cells at 14 dpt stained with SATB2 and CTIP2. Scale bar: 400 um. Quantitation of layering: distance of each cell from the outside border of the graft. N = 3 mice. D. Sections showing areas in E-H. Scale bars: 1 mm. E-H. Neuronal projections from transplanted cells stained for GFP (n = 3/3 for each area). Scale bar: 50 μm I. Host striatal response to stimulation of ChR+ axons. Raster plot of train stimulation. 1 ms pulse from 100 um optrode with 4-6 mW light was used for the representative trace. N = 3 mice. J. Line scan using 2-photon microscopy to measure changes in GCaMP6f fluorescence in response to electrical and optical stimulation with and without synaptic blockers NBQX and D-APV. Graph of cell response in blue and green. Response in the presence of blockers in red. Bottom right: quantification. N = 3 mice, n = 6 cells. DL, deep layers; HIP, hippocampus; NCX, neocortex; UL, upper layers.

By comparing the neuronal projections described in the Allen Brain Institute database for neurons located in the same primary somatosensory cortical region as our transplants, we found that at 14 dpt graft-derived mGFP+ projections appropriately targeted the motor cortex, striatum, and surrounding somatosensory cortex, but not inappropriate targets such as the thalamus (Figure 1D-H, n = 3 out of 3 grafts targeting motor cortex, striatum, somatosensory cortex, and n = 0 out of 3 targeting thalamus). To test if donor neurons could synaptically activate host neurons, we transplanted donor cells from E12.5 neocortices carrying the *Foxg1^Cre^* allele and a conditional channelrhodopsin (ChR2) allele, enabling the activation of transplanted neurons with light. One month after transplantation, in awake freely moving recipient mice, we recorded the activity of host striatal neurons in response to stimulation of donor-derived axons using an electrode affixed to a fiberoptic that delivered 1 msec pulses of 2 milliwatt blue light. Host neuron responses followed a train of light pulses used to stimulate donor fibers at 20 hertz, indicating that transplanted neurons were synaptically communicating to host neurons (Figure 1I).

Conversely, to examine if donor neurons respond to input from host neurons, we transplanted donor cells from E12.5 neocortices carrying the *Foxg1^Cre^* allele with a conditional GCaMP6f allele, which encodes a calcium indicator, into the adult brains of mice in which host neurons expressed ChR2. Using acutely prepared brain slices from mice harboring 2-month-old grafts, we stimulated host neurons both electrically and optically at 200 μm from the graft edge and measured calcium transients. Both electrical and optogenetic stimulation produced responses in the transplanted cells that were blocked by antagonists of glutamatergic synaptic transmission, D-APV and NBQX, which target NMDA and AMPA receptors, respectively (Figure 1J). Our findings demonstrate that transplants of dissociated fetal cells at stroke sites generate healthy, well-differentiated neurons that can synaptically integrate with the host.

### Dissociated donor endothelial cells form vessels within grafts and circulate blood

Given that we observed successful engraftment and that donor vascular cells were included in the donor cell population, we evaluated the role of vascularizaton in the engraftment process. If the inclusion of donor vascular cells in a transplanted cell population contributed to graft survival, then we might expect donor-derived vascular cells to form vessels that anastomose with host vessels to circulate blood within the graft. In addition to the presence of mature neurons within our grafts, we observed extensive blood vessels in all grafts examined (>50 mice) (Figure 2A). To determine if the vessels in the grafts were donor and/or host derived, we transplanted cells from E12.5 embryos carrying an eGFP reporter driven by *Mesp1^Cre^* to label cells of the mesoderm lineage, which in the brain consists primarily of vascular cells (Supplementary Figure 1). To control for potential variations in donor cell preparations and host immune responses, donor cells were transplanted into both hemispheres of adult wild type mice that had sustained a unilateral left dMCAO. Transplants were performed at 3 and 7 days post-stroke, times corresponding to the subacute phase when transplants might be most clinically relevant (Carmichael, 2016a, 2016b; Doyle et al., 2008).

**Figure 2.**
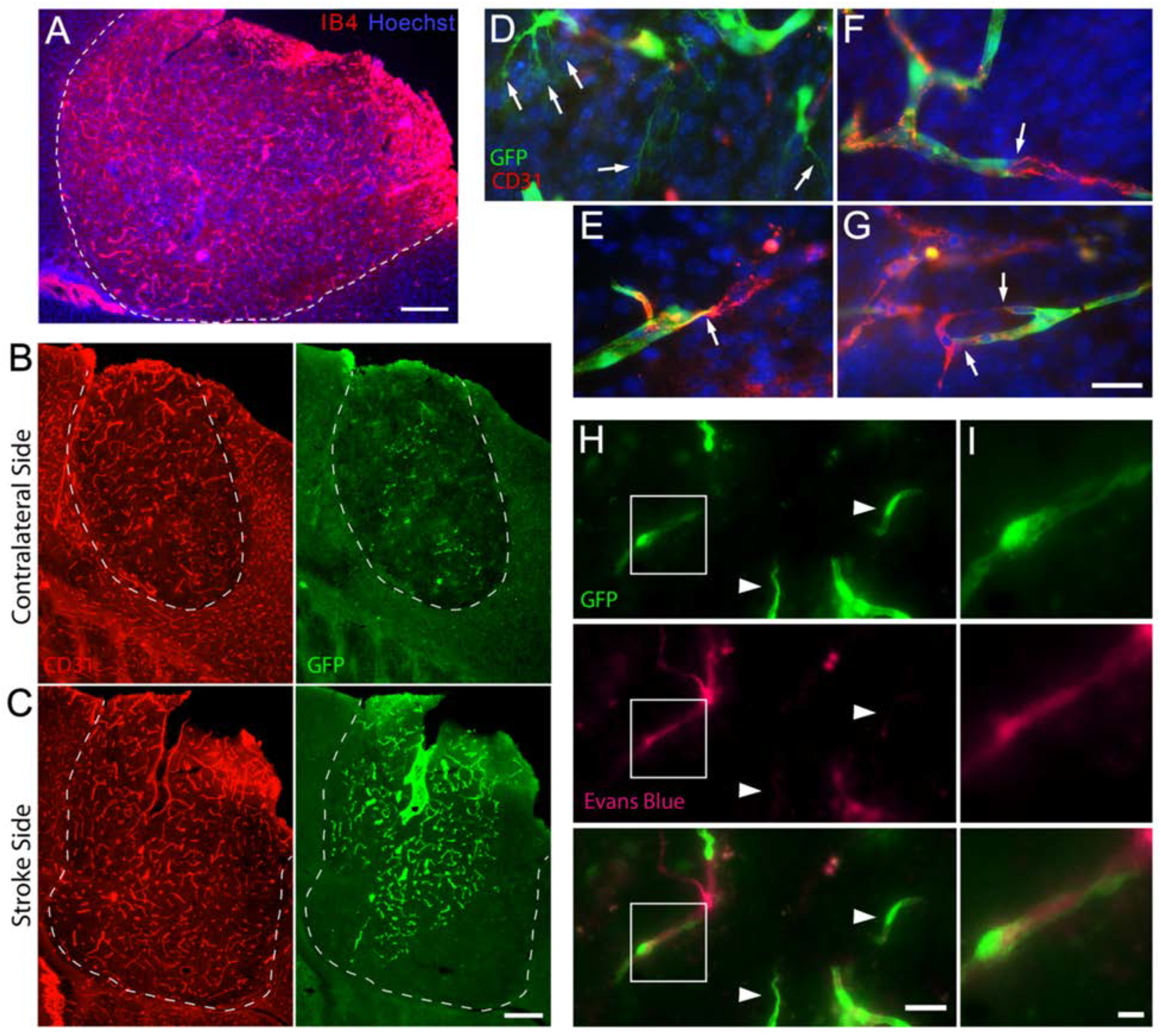
Transplants are vascularized with both host and donor-derived endothelial cells. A. Transplant site stained with endothelial cell marker IB4 and nuclear marker Hoechst. Scale bar, 200 um. B, C. Transplant on contralateral side and stroke side (C) of the same mouse with GFP labeled donor vascular cells and stained with endothelial marker CD31. Scale bar: 200 μm. D - G. Representative examples of fusion of host (red) and donor (green) vessels; filopodial sprouting (arrows, D), contact (arrow, E), apical membrane insertion (arrow, F), and completion of fusion (arrows, G). Scale bar, 20 μm. H. Donor derived vessels (GFP+) carrying Evans Blue tracer (red). Immature-looking vessels that have not yet anastomosed do not contain Evans Blue (arrowheads). Scale bar, 20 μm. I. Higher magnification of boxed regions. Scale bar, 5 μm.

Two weeks after transplantation, grafts at the stroke site and contralateral site contained both eGFP-labeled donor- and host-derived vasculature (Figure 2B,C and see also quantification in Figure 3), demonstrating that transplanted vascular cells maintain their ability to form vessels after transplantation. Due to the general health and large size of the grafts, we hypothesized that the donor-derived vessels were functional, which would require anastomosis between eGFP+ microvessels and host vessels. In support of this hypothesis, we observed morphological evidence for several of the steps involved in anastomosis (Lenard et al., 2013). Donor endothelial tip cells exhibited sprouting, contact, apical membrane insertion, and fusion with host vasculature (Figure 2D-G) and host vascular cells, in turn, exhibited morphologies suggestive of reaching out and fusing with donor vessels (Supplementary Figure 2A). Consistent with a role for microglia in anastomosis (Arnold and Betsholtz, 2013; Fantin et al., 2010), we also observed microglia, which were both host and donor-derived, closely associated with tip cells (Supplementary Figure 2B-E).

**Figure 3.**
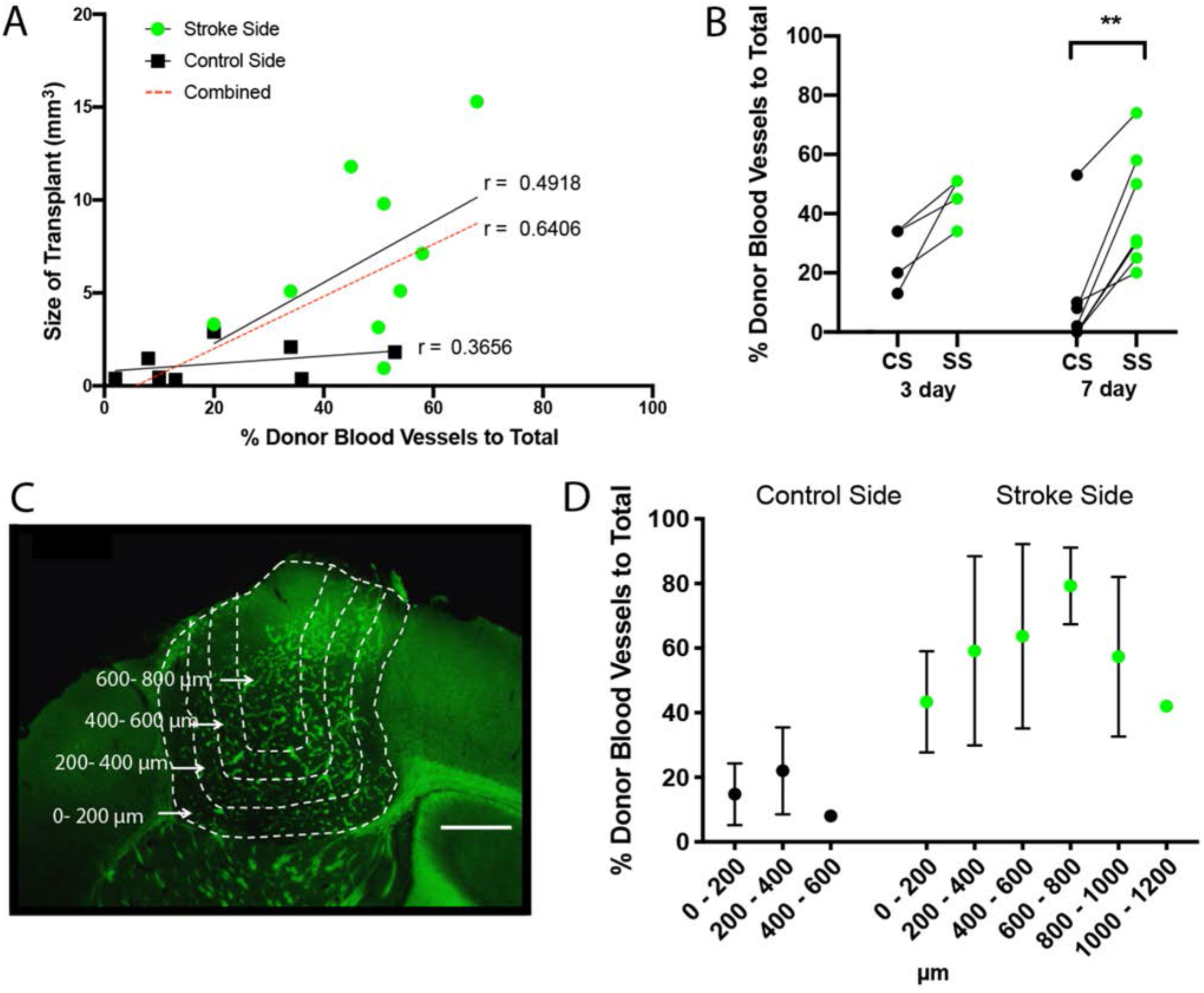
Grafts at stroke sites were larger with more donor-derived vessels than at nonstroke sites. A. Size of grafts on both the contralateral side (black), stroke side (green), and combined (red dotted line) were plotted against the percent contribution of donor vessels within the graft. Pearson’s correlation coefficient. B. Percent contribution of donor vessels compared pairwise between the contralateral and stroke sides for each host brain. CS, control side; SS, stroke side. Wilcoxon matched-pairs signed rank test **p = .033. C. Representative schematic of regions that were analyzed for quantification in (D). Scale bar, 400 μm. D. Percent contribution of donor derived vessels within each region of both stroke and contralateral side grafts.

To address whether that donor-derived vasculature circulates blood, we injected host mice intraperitoneally with Evans Blue, which enters the circulatory system. This dye was then detected within the eGFP+ vessels in grafts on both the stroke-affected and the contralateral sides, demonstrating that donor-derived vasculature circulates blood (Figure 2H, n=3/3 mice). These findings support the possibility that donor cell-derived vessels are essential to improving neocortical engraftment.

### Grafts at infarct sites have more donor-derived vessels and are larger than at control sites

Two weeks after transplantation, grafts at stroke sites were significantly larger than those in the contralateral hemisphere (Figure 3A). The large grafts had a greater cell number than initially injected (100,000 cells injected reaching 1,300,000 +/- 930,000 cells at 2 weeks, N = 9), indicating extensive cell proliferation. These results suggest that the post-stroke environment is well suited for cell engrafting.

If our hypothesis is correct that donor-derived vessels are required to support large grafts, then only grafts with greater contributions of donor-versus host-derived vasculature should be large. To address this possibility, we performed transplants at stroke sites and control contralateral sites. In the days following dMCAO, host angiogenesis peaks at 3-4 days after ischemia, followed by a decrease at about 7 days (Hayashi et al., 2003). Therefore, we performed transplants at 3 and 7 days following dMCAO to control for the changing ischemic environment, and quantified donor and host vessels within grafts 2 weeks later using AngioTool software (Zudaire et al., 2011). We found that the percentage of donor-derived vessels was significantly greater on the stroke side, which had larger grafts, compared to the control side for the 7-day group and trending for the 3-day group (Figure 3B). In addition, and consistent with our hypothesis, the fraction of donor-derived vessels correlated with graft size when the data for the stroke and contralateral sides were analyzed separately or together (Figure 3A). These results suggest not only that the post-ischemic environment promotes graft growth, but also that donor vascular cells may be required to generate and sustain grafts beyond a certain size.

### Donor-derived vasculature is found increasingly toward the center of grafts

If indeed donor-derived vessels allow transplanted cells to form larger grafts, then we would expect that the ratio of donor- to host-derived vessels should be greater toward the center region of the transplant where demand for oxygen and nutrients might be greatest and where host vessels would take the longest to reach. To test this hypothesis, we again used donor cells from *Mesp1^Cre/+^;eGFP-reporter* embryos to visualize donor-derived vasculature within grafts sites at control and stroke sites. We calculated the percent of donor vessels by quantifying the ratio of donor-derived eGFP+ vessels to total CD31+ vessels in nested 200 μm bins from the edge of the graft-host border to the center of the graft (Figure 3C). Accordingly, the relative contribution of donor-derived vessels appeared to increase up to 800 μm from the graft-host border (Figure 3D). These data further support our hypothesis that inclusion of dissociated donor vascular cells promotes vessel formation necessary to sustain larger grafts.

### Removing vascular cells from the donor cell population results in decreased graft size

To determine if donor vascular cells are required to achieve larger grafts, we initially attempted to subtract them using two methods. First, CD31+ vascular endothelial cells were eliminated with immunobeads by magnetic activated cell sorting (MACS), and second, GFP+ vascular endothelial cells, whose fluorescence is activated by *Mesp1^Cre^*, were removed with fluorescence activated cell sorting (FACS). However, both purification processes greatly reduced overall cell viability, as shown in control experiments in which cells that underwent the same sorting treatments but without removal of vascular cells resulted in much smaller grafts (Supplementary Figure 3). For FACS, reduced overall viability of donor cells occurred despite minimizing stress on cells by using a minimal flow rate and a maximal nozzle diameter.

Therefore, we took an alternative approach to depleting vascular cells from the donor cell population. Two main cellular sources provide blood vessels to the developing neocortex: the perineural vascular plexus (PNVP) and the periventricular plexus (PVP). The PNVP is contained within the nascent meninges by E9, while the PVP populates the cortex approximately two days later at E11.5 (Vasudevan et al., 2008). These two plexuses then fuse to complete the vascular network. At E12.5, the majority of vascular cells remain contained within the pia (Supplementary Figure 1), and their surgical removal drastically diminishes the percentage of donor-derived vascular cells within our transplants (48 ±14% versus 2.7 ±3.9%, Figure 4A,B). Thus, to determine if donor vascular cells are required to achieve larger grafts, we depleted them from the donor cell population by removing the pia before dissociating E12.5 donor cortical tissue. We performed the transplants in stroke-affected and contralateral control sides, with a one-week delay between dMCAO and transplantation. The resulting 2-week-old grafts exhibited similar fractions of SATB2+, CTIP2+, and total neuronal cells, although the grafts without pia appeared less pseudo-layered (Supplementary Figure 4).

**Figure 4.**
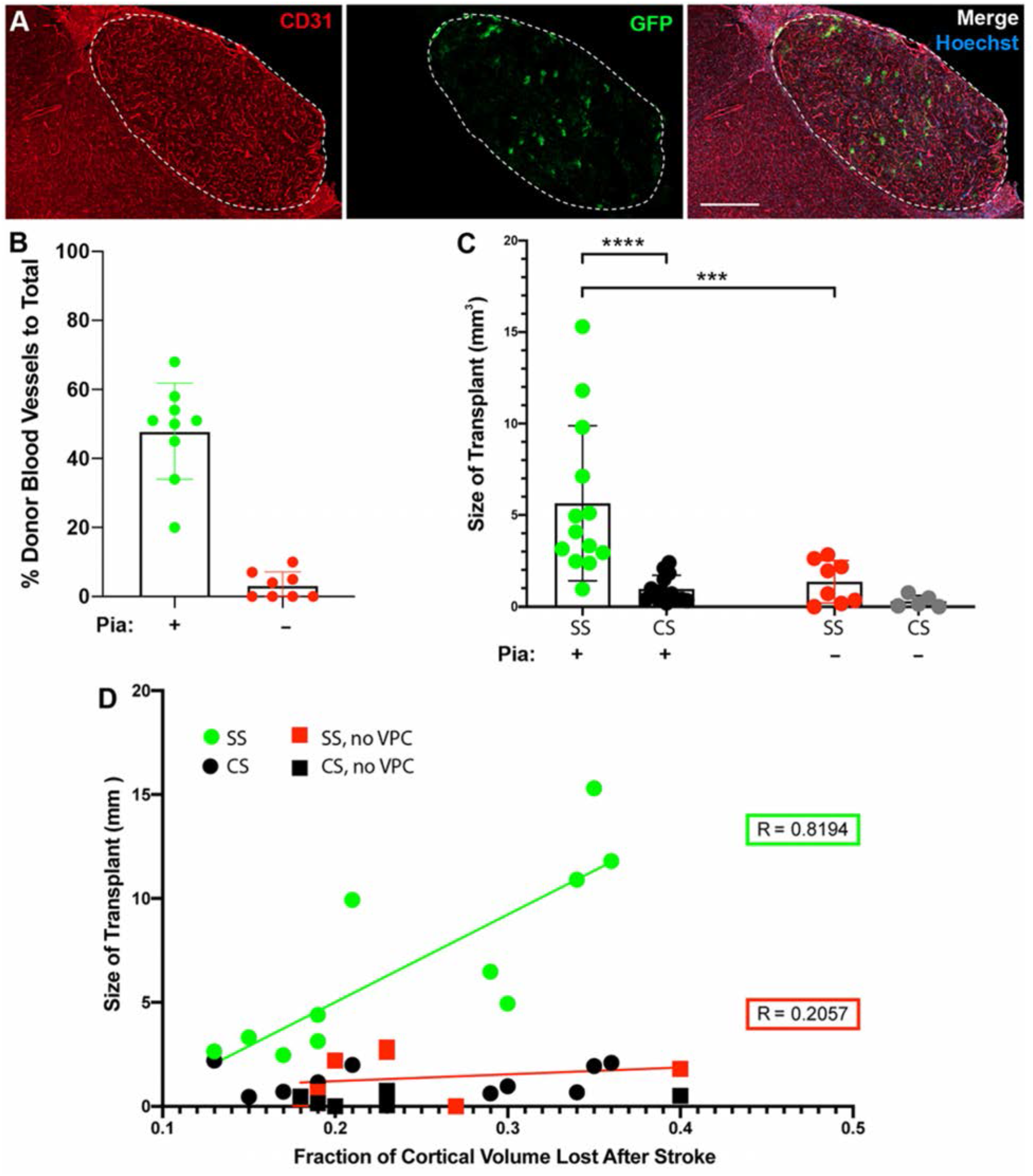
Removal of donor-derived vessels results in smaller grafts. A. Representative transplant site from a donor population of cells that excluded pia, stained for CD31 (total vessels, red), donor-derived vessels (GFP, green), and Hoescht (blue). Scale bar: 500 μm. B. Percent contribution of donor-derived vessels with (N = 9) and without (N = 8) pial cells in the donor cell population at stroke sites. Data are presented as Mean ± s.d. Mann-Whitney test, ****p <.0001. C. Size of both stroke and contralateral side grafts with (green) and without (red) donor derived vessels. CS, control side; SS, stroke side. Data are presented as Mean ± s.d. Mann-Whitney test, ***p < 0.0002, ****p < 0.0001. N = 8 (+pia), N = 5 (-pia). D. Correlation between the extent of stroke, measured as the lost fraction of cortical volume normalized to the contralateral hemisphere, and the size of transplants. With pia (green) N = 11; without pia (red) N = 8, Pearson’s correlation. CS, control side; SS, stroke side; VPC, vascular precursor cells.

Consistent with our hypothesis, the average size of the grafts at stroke sites decreased from 5.64 ±4.3 mm^3^ for donor populations containing dissociated pia to 1.34 ±1.16 mm^3^ for donor populations without pia (Figure 4C). Moreover, despite a correlation between stroke size and graft size when vascular cells are included in the donor cell population, when vascular cells were depleted graft size remains small regardless of stroke size (Figure 4D). These findings indicate that the depletion of vascular cells from the donor population significantly limits the size of grafts.

## Discussion

The repair of neocortical tissue by transplantation of neural cells will require several methodological improvements, beginning with increased survival of donor cells. Although the inclusion of vascular cells in neural cell transplants was hypothesized to promote graft survival and integration, clear evidence confirming this hypothesis was lacking. Our study shows that including blood vessel-forming progenitors in a transplant cell population is necessary to surpass a nominal graft size in the mouse neocortex. First, The large grafts contained upper and lower layer neuronal subtypes that synaptically connected with appropriate targets, highlighting general graft health. Second, grafts with greater contributions of donor-derived vessels were larger. Third, the contribution of donor-derived vessels appeared to increase as a function of distance from the graft-host border. Finally, whereas grafts with donor vascular cells reached 15 mm^3^, excluding vascular cells from the donor cell population limited grafts to a maximal size of 3 mm^3^.

Obtaining large grafts from dissociated cells has been challenging. NSCs derived from cell lines are inherently comprised of cells from a pure neuronal lineage and lack glial and vascular progenitor cells. Among neocortical transplantation studies that use single-cell suspensions, few have shown that new neurons were generated in substantial numbers. Too few surviving transplant-derived neurons would likely limit their connectivity with the host and their contribution to circuitry that underlies behavior. The results of our present study showing growth and survival of grafts derived from a mix of neural and vascular cells suggests that poor outcomes obtained using purified cell populations are likely due at least in part to the lack of support from vessel-forming vascular cells. Our findings here suggest that the vessels generated from donor cells supply the developing graft, particularly towards its center, with necessary oxygen and nutrients more quickly than would ingrowing host vessels. Notably, the donor-derived vessels self-organized from single-cells into an extensive vasculature, a property which may be advantageous in clinical settings, where individual cell types would be grown separately in culture prior to mixing them for transplantation.

To highlight the importance of including vascular cells in the transplant cell population, we removed the pia, the major source of vascular cells, prior to dissociating donor neocortical tissue. In addition to providing vasculature to the parenchyma, the pia may have other functions in neuronal differentiation (Siegenthaler and Pleasure, 2011), leaving open the possibility that pia-derived factors other than vasculature may contribute to successful engraftment. Similarly, given that microglia are known to participate in vascularization and anastomosis (Arnold and Betsholtz, 2013; Fantin et al., 2010) and that both donor and host microglia are present in grafts (Supplementary Figure 2), the extent to which donor-derived microglia might promote graft vascularization and health remains unclear. Future experiments will be required to determine the benefits, if any, of including microglia in neocortical cell transplants.

In this study, we used stroke as a physiologically relevant injury model. Notably, grafts at stroke sites were larger (with a higher percentage of donor-derived vessels) than grafts at nonstroke sites. This suggests that the environment in which cells are transplanted also plays an important role in graft vascularization, survival, and growth, consistent with observed dynamic post-stroke revascularization (Williamson et al., 2020). Several possible factors could account for the better vascularization and engraftment at stroke sites. Hypoxia, angiogenic factors such as VEGF, and inflammation are elevated at stroke sites for up to three weeks and could promote vessel formation by the donor vascular endothelial cells (Hayashi et al., 2003; Krock et al., 2011).

In this study we did not examine the behavioral consequences of the grafts for three reasons. First, our current grafting method is not likely to yield functional neocortical tissue due to their abnormal cytoarchitecture (despite pseudo-layering, which does not align with host layers) and cell composition (few if any interneurons for example are present in E12.5 donor tissue). Second, the mouse neocortex compensates relatively quickly and efficiently for loss of cortical tissue, making putative minor contributions to improved behavior difficult at best to detect. Third, simply examining behavior does not distinguish between well-documented bystander effects (Zhang et al., 2020) and actual repair (whereby neuronal activity of the graft itself provides behavioral benefits). Future advances will be required before neocortical grafts can be shown to function as normal neocortical tissue.

Nevertheless, the more that new graft tissue recapitulates the endogenous makeup of neocortical cell types, the better its chances will be of providing a new substrate for lost function. The neocortical cell population is composed of less than 50% neurons (von Bartheld et al., 2016), with the rest of the cell types each being vital for the health and function of neurons. Prime among these cell types are vascular cells that can rapidly form vessels and anastomose with the host to deliver oxygen and nutrients to the graft more quickly than would ingrowing host vessels. This is particularly relevant to ischemic strokes in humans where lesions, although varying greatly in size, have an average volume for tissue loss of roughly 54,000 mm^3^ (or 54 ml) (Saver, 2006). Hence, transplanted cells used to replace even a fraction of lost tissue will likely depend on the inclusion of blood vessel-forming cells. Our results show that transplanting vascular progenitor cells with neural precursors supports increased graft size. This combined cell transplant approach will provide a platform on which to begin reassembling other cell types in future studies aimed at optimizing the ability of grafts to repair damaged brain tissue.

## Experimental Procedures

### Animals

Animal handling followed an approved protocol by the Albert Einstein College of Medicine Institutional Animal Care and Use Committee in accordance with National Institute of Health guidelines. CD1 adult male and female mice of 2-5 months of age were used as hosts in this study. Transgenic donor embryos were generated from mice of mixed backgrounds. For experiments analyzing donor vascular cell contribution, males carrying *Mesp1^Cre/+^* were crossed to females homozygous for *Rosa26CAG^lox-stop-lox-eGFP^* or *Rosa26^lox-stop-lox-tdTomato^* (Saga et al., 1999; Sousa et al., 2009). Males with *Foxg1^Cre/+^* were crossed with females homozygous for *Rosa26^lox-stop-lox-GCaMP6f^* or *Rosa26^lox-stop-lox-ChR2-YFP^* (Hébert and McConnell, 2000; Madisen et al., 2012; Madisen et al., 2015) for electrophysiology experiments. Mice were housed on a 12 hour light-dark cycle with a maximum of five animals to a cage and provided with water and chow ad libitum.

### Induction of Distal Middle Cerebral Artery Occlusion (dMCAO)

Mice were anesthetized under 1.5% isoflurane and after a small incision was made in the skin and a skin flap folded back, a small hole was drilled in the skull over the distal left middle cerebral artery (MCA). Permanent cortical ischemia was induced by cauterization and disconnection of the MCA at a location in the distal trunk before splitting into the anterior and posterior branches. The skin was then sutured closed.

### Preparation of Donor Cells from Mouse Embryos and Transplantation

Embryos of the desired genotype were harvested from pregnant females at E12.5. The developing cortex was collected with overlying primitive leptomeninges (arachnoid and pia mater), except for the experiments shown in Fig. 4, where the leptomeninges were removed for the experimental group of transplants. Cortices were incubated in accutase for 20 minutes to ensure dissociation of cells. The single cell nature of the donor cells was confirmed under light-field microscopy. After spinning, cells were resuspended at 50,000 cells/μl in medium containing Neurobasal, N2, B27, penicillin-streptomycin, GlutaMax and HEPES buffer. Host mice that had undergone dMCAO 1, 3, and 7 days prior were anesthetized with 1.5% isoflurane and injected with 2 μl of cell suspension bilaterally at −0.5 mm posterior and +/- 3.3 mm medio-lateral to bregma, at a depth of 2 mm from the surface of the skull, which corresponds to the primary somatosensory cortex. Cell suspensions were injected using a 30 gauge Hamilton syringe at a rate of 0.150 μl/min. The needle was left in for 5 minutes before slowly retracting to minimize backflow.

### Magnetic Activated Cell Sorting

MACS was performed in accordance with the manufacturer’s protocol (MACS Miltenyi Biotec Order No. 130-097-418). In brief, cortical cells were isolated from embryos as stated above and passed through a 40 μm filter. Cells were resuspended in buffer and incubated with CD31 MicroBeads for 15 minutes at 4°C. After washing in buffer, the cell suspension was passed through a magnetic field in MACS column that was affixed to a MACS Separator. Flow through of unlabeled cells was collected and resuspended in transplantation (above) and immediately injected into host mice.

### Immunofluorescence

Mice were perfused with 4% paraformaldehyde (PFA) intracardially 2 weeks after transplantation (unless otherwise specified) and post-fixed overnight in 4% PFA at 4°C. Brains were cryoprotected with 30% sucrose for 48 hours, embedded in OCT, and stored at −80°C. Sections were cut at 30 μm and immunostained as floating sections. Where needed, antigen retrieval was performed by heating sections for 45 minutes at 90°C in 10 mM sodium citrate pH 6.0. The following primary antibodies were used: CD105 (1:100)) Cat No. 120402 BioLegend, PECAM (1:100) Cat No. ab28364 AbCam, GFAP (1:500) Cat No. 13-0300 ThermoFisher, GFP (1:250) Cat No. A-11122 ThermoFisher, Satb2 (1:500) Cat No. ab92446 AbCam, Ctip2 (1:500) Cat No. ab18465 AbCam, Ki67 (1:500) Cat No. ab15580 AbCam, Iba1 (1:500) Cat No. 019-19741 Wako. Primary antibodies were incubated overnight at 4°C and in Alexa-488, Alexa-568 and Alexa-647 secondaries (1:500 Invitrogen) for 1 h at room temperature.

### Image Analysis of Transplants

Blood vessels were quantified using the software AngioTool (Zudaire et al 2011). The area of donor-derived blood vessels was divided by the area of total blood vessels to determine the fraction of vessels stemming from the donor population. Size of transplants was quantified using ZEN software (Blue edition, Zeiss).

### Acute Cortical Slice Preparation

Mice used for GCaMP6f imaging were sacrificed two months after transplantation. Mice were anesthetized using 4% Isoflurane and transcardially perfused with 20 mL of cold NMDG solution containing in (mM): 93 NMDG, 2.5 KCl, 1.25 NaH_2_PO_4_, 30 NaHCO_3_, 20 HEPES, 25 glucose, 5 sodium ascorbate, 2 Thiourea, 3 sodium pyruvate, 10 MgCl_2_, 0.5 CaCl_2_, brought to pH 7.35 with HCl. Mice were decapitated and their brains were carefully removed for coronal slice preparation (350 μm thick) in cold NMDG solution using a VT1200s microslicer (Leica Microsystems Co.). Acute slices were collected and transferred to a chamber filled with artificial cerebrospinal fluid (ACSF) solution containing (in mM): 124 NaCl, 2.5 KCl, 26 NaHCO_3_, 1 NaH_2_PO_4_, 2.5 CaCl_2_, 1.3 MgSO_4_ and 10 glucose. The chamber was kept in a warm-water bath at 33-34°C during slicing and then at room temperature until recordings started. All solutions were equilibrated with 95% O_2_ and 5% CO_2_ (pH 7.4).

### Two-photon Calcium Imaging and Optogenetics in Slices

Acute coronal slices were brought to a submersion-type recording chamber perfused at 2 ml/min with ACSF supplemented with the GABA-A receptor antagonist picrotoxin (100 μM) and maintained at 28 *±* 1 °C. The transplantation site was identified using an epifluoresence lamp with excitation/emission settings for GFP visualization. For electric stimulation, a monopolar stimulation glass electrode was placed ~200 μm from the transplantation site using differential interference contrast (DIC) microscopy. The Invitro Ultima 2P microscope (Bruker Corp) designed for acute slice imaging contains an Insight Deep See laser path that was tuned to 920-945 nm to acquire GCaMP6f signals via the “green” photomultiplier tube (PMT) using minimal pockel power at the 60x objective (Nikon, 1.0 NA). GFP positive cell bodies were identified in the field of view using 512 x 512 pixel resolution and line scan analysis software was implemented to designate a region of interest (ROI) to capture calcium transients (CaTs). To elicit CaTs, 10-20 electrical pulses at 10 ms inter-stimuli-interval using square wave pulse widths of 100 μs duration were delivered using an Isoflex (A.M.P.I.) stimulator. IgorPro 6 custom software was used to trigger a Master 9 pulse stimulator (A.M.P.I.) to induce electrical stimulation and image acquisition software. For each ROI, three-trials were obtained with a baseline fluorescence duration of 400 ms followed by electrical stimulation and CaT induction. After CaT acquisition trials were completed a drug cocktail containing NBQX (10 μM) and D-APV (25 μM) was washed-in for at least 10 min and CaT acquisition trials were repeated. For optogenetic stimulation in Channelrhodopsin (ChR2) host animals the Invitro Ultima 2P microscope contains a Coherent 473 nm laser path. The optical stimulation was defined using customized Mark Point software (Bruker Corp) where 10 stimulation points surrounded a responsive ROI previously identified with electrical stimulation. The Mark Point software was set to deliver 1 optical pulse at 20 ms inter-stimulus-interval (4-8 mW, 1-2 ms pulse duration) at each stimulation point resulting in a total of 10 optical pulses. Upon optical stimulation a high-speed shutter blocked the light path of the “green” PMT preventing image acquisition until optical stimulation terminated. When optical stimulation was complete, the shutter retracted and image acquisition resumed capturing the remaining profile of the CaT. Three imaging trials were conducted followed by NBQX and D-APV wash-in as previously described.

### Calcium Transient Analysis and Statistics

CaTs were acquired with a 1-2 min inter-trial-interval and analyzed. The ΔF/F was calculated for a 200 msec time-bin before and after NBQX and D-APV application, and it was compared across ROIs for both electrical and optical stimulation. In CaTs elicited by optical stimulation, the 200 msec time-bin was defined as the time after the high-speed shutter retracted and maximal CaT elevation was achieved. All peak ΔF/F values were tested for normal distribution using the Shapiro-Wilk test and p value was set to 0.05 for statistical significance. A paired *t*-test was used for ΔF/F values before and after drug treatment when data was normally distributed and sample size was > 6 slices. A paired sample Wilcoxon Signed Rank test was used for statistical significance when data was not normally distributed.

### In Vivo Optogenetic Stimulation of Transplant Fibers

Fibers from the transplant were stimulated optogenetically with a 447 nm wavelength laser (OEM Laser Systems, Inc.) in awake head-fixed mice using a custom-made optrode. Optrodes contained a 100 μm optical fiber (Multimode Fiber, 0.22 NA, High-OH 105 μm Core, ThorLabs, FG105UCA) that was glued to a tungsten electrode (WPI, TM33B20, shaft diameter = 0.256 mm) with epoxy (Devcon – 5 Minute Epoxy). For post-mortem identification of the optrode location, the optrode was coated with DiI prior to placement in the striatum (Invitrogen, V22889). Coordinates ranged from −3.8 to −4.6 mm, AP; ±1.2 mm ML; −4.5 mm below DV. Once the optrode contacted the surface of the brain, the well around the recording site was filled with sterile saline, and the reference wire was placed in the saline. The optrode was then lowered to the targeted depth. For the optical single-pulse stimulation protocol, 1 ms pulses of light were delivered at five-second intervals. For the train stimulation protocol, five light pulses of 1 ms duration were delivered at 20 Hz with an interstimulus interval of 10 seconds. For all protocols, at least 40 trials were repeated for each cell. Isolated extracellular single-unit signals were amplified 2,000x using a custom-made differential amplifier (1 Hz to 10 kHz, RC band-pass filter) and digitized at 20 kHz (National Instruments, PCI6052E). Data were collected using custom written software in LabVIEW (National Instruments), and single units were sorted offline according to principal component analysis (PCA) using Plexon® Offline Sorter™. Data were analyzed with custom-written programs in MATLAB10 (Mathworks). A peri-stimulus time histogram (PSTH) using 2 ms bins was made for each cell.

### Statistical Methods

Statistical analyses were performed using GraphPad Prism. Data were analyzed for statistical significance using Wilcoxon Ranked test for pairwise analyses of cells and Mann-Whitney U test for comparisons between two groups. Sample sizes and p values for each experiment are listed in the figure legends.

## Acknowledgements

We are grateful for grant support from the SENS Research Foundation (JMH), New York State Department of Health NYSTEM Program for shared facility grant support C029154 (JMH), NYSTEM C34875GG (JMH), NYSTEM Einstein Training Program in Stem Cell Research C30292GG (JK and MGP), Brain Research Foundation (JMH), NIH NS088943 (JMH) and MH125772 (PEC), Ruth L. Kirschstein NRSA Fellowship F31MH109267 (PJL), as well as seed funding from the Albert Einstein College of Medicine. We are also grateful to lab members for feedback on the manuscript.

## Author Contributions

JK designed and performed most of the experiments and wrote the manuscript, JY performed early dMCAOs, HG performed the initial neuronal analyses, BG performed initial analyses of vascularization in grafts, SK performed early ChR2 optogenetic experiments, KA and PL performed the GCaMP6f analyses, MGP established the initial transplant paradigm, YDT contributed quantifications of percent donor vessels in grafts, KM analyzed embryonic tissue, RSZ provided guidance for the dMCAO procedure, DAW provided guidance with vascular analyses, PEC supervised the GCaMP6f experiments and edited the manuscript, KK supervised all aspects of the study and edited the manuscript, JMH supervised all aspects of the study and edited the manuscript.

## Supplemental Information

**Supplementary Figure 1.**
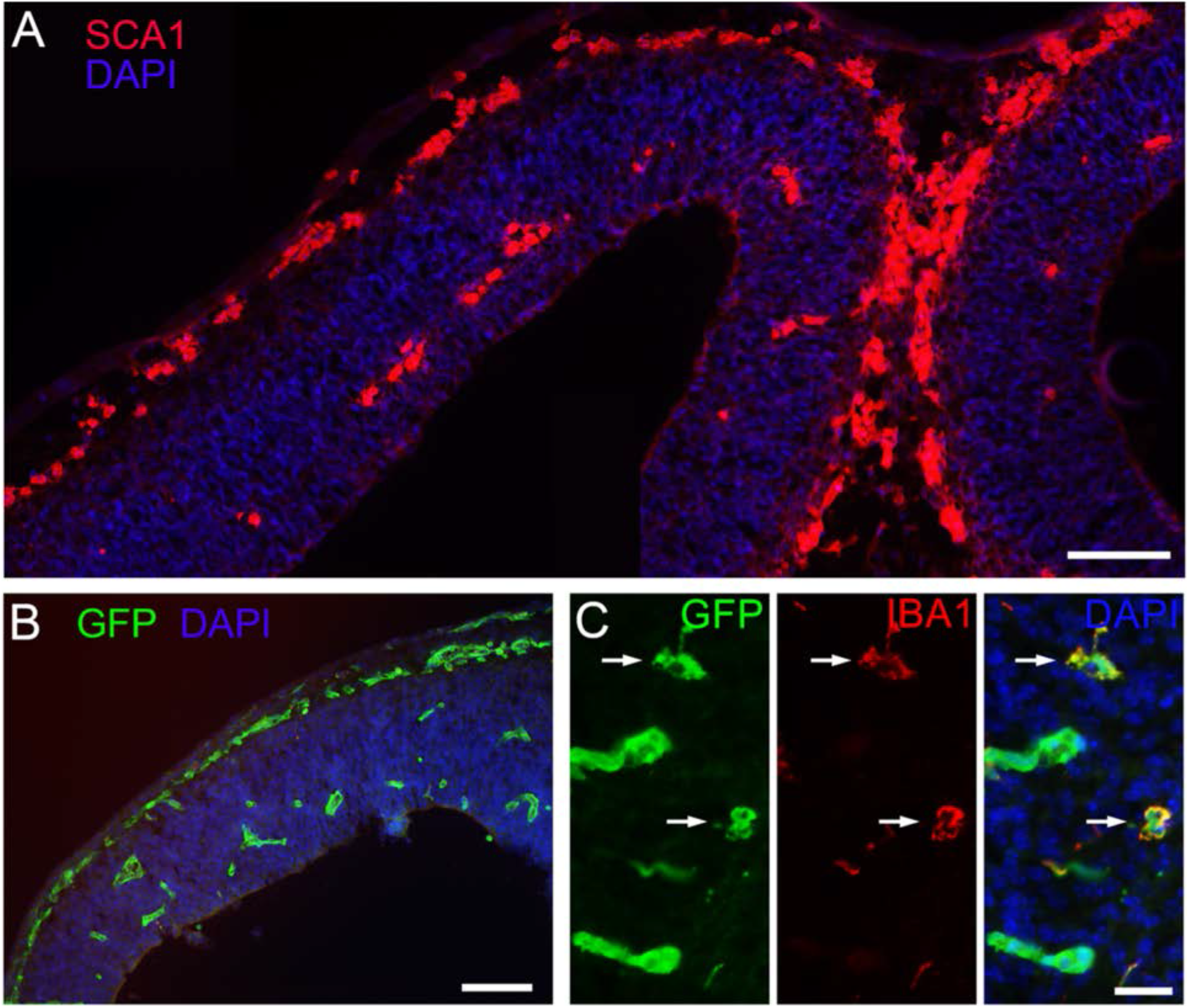
The E12.5 developing mouse neocortex is comprised of vascular and microglial precursor cells in addition to the neuroepithelium. **A)** SCA1 immunostaining marks vascular cells. **B)** Embryos carrying *Mesp1^Cre/+^;Rosa26^CAG-loxSTOPlox-eGFP^* express GFP in mesodermal derivatives, primarily vascular cells in the brain, but also microglial precursors. **C)** Microglial precursors are labeled with GFP and IBA1 (arrows) in E12.5 mouse neocortex. N = 4/4 mice for each immunostain. Scale bars, A,B 50 μm; C, 15 μm.

**Supplementary Figure 2.**
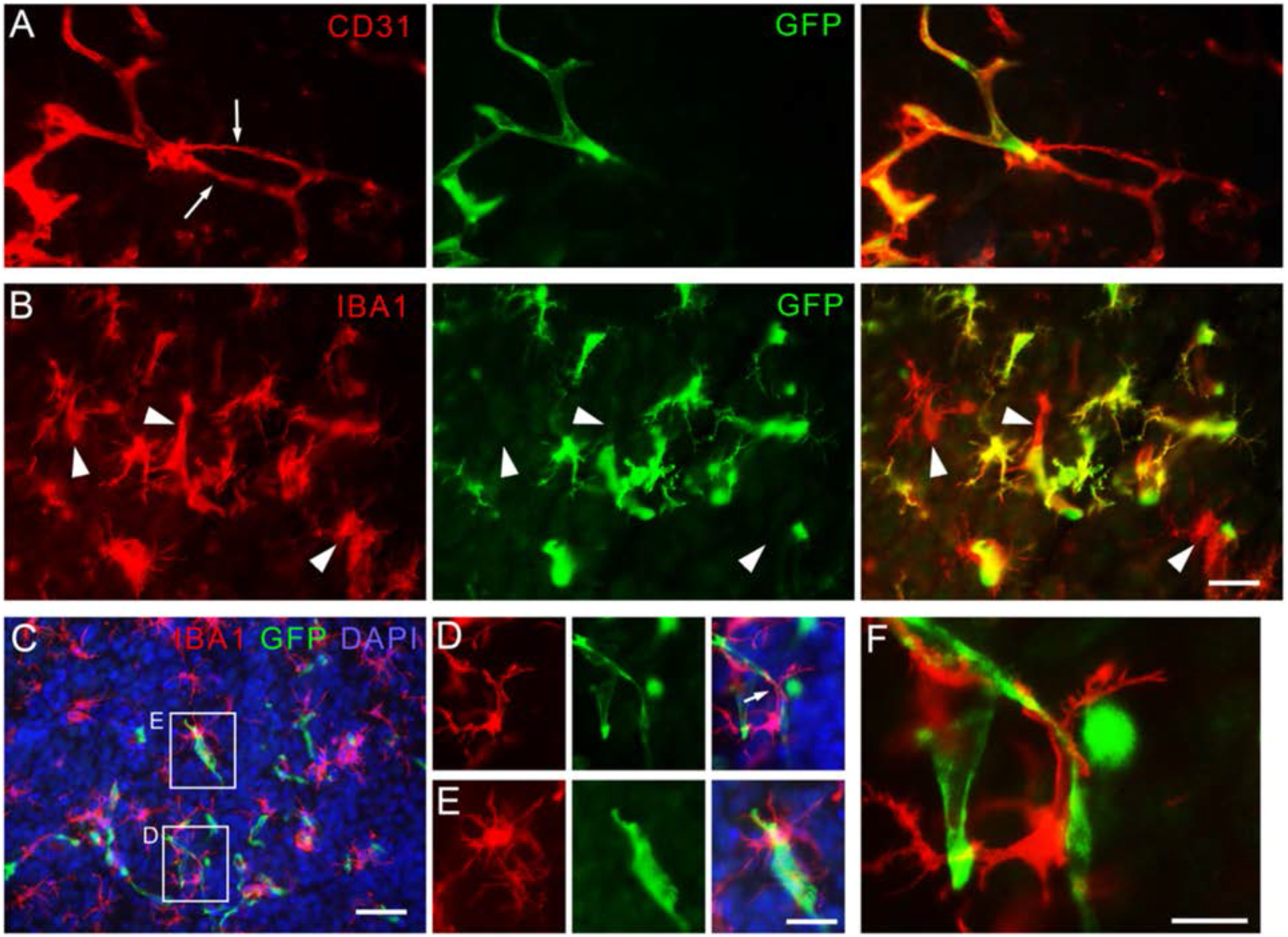
Host vascular cells send processes to donor-derived vessels, which are associated with microglia. **A)** Example of thin PECAM+ host (red only) vascular processes (arrows) making contact with a donor-derived GFP+ vessel (green + red) in a graft. N=8/8 grafts. **B)** IBA1+ microglia within grafts are both donor-derived (green + red) and host-derived (red). N=6/6 grafts. **C)** Microglia are associated with budding vessels. **D)** and **E)** are higher magnification views of boxed areas in C). **F)** Higher magnification of area indicated by an arrow in D) showing an example of a microglial process associated with a vascular endothelial process. Scale bars: A,B,D,E, 20 μm; C, 40 μm; F, 10 μm.

**Supplementary Figure 3.**
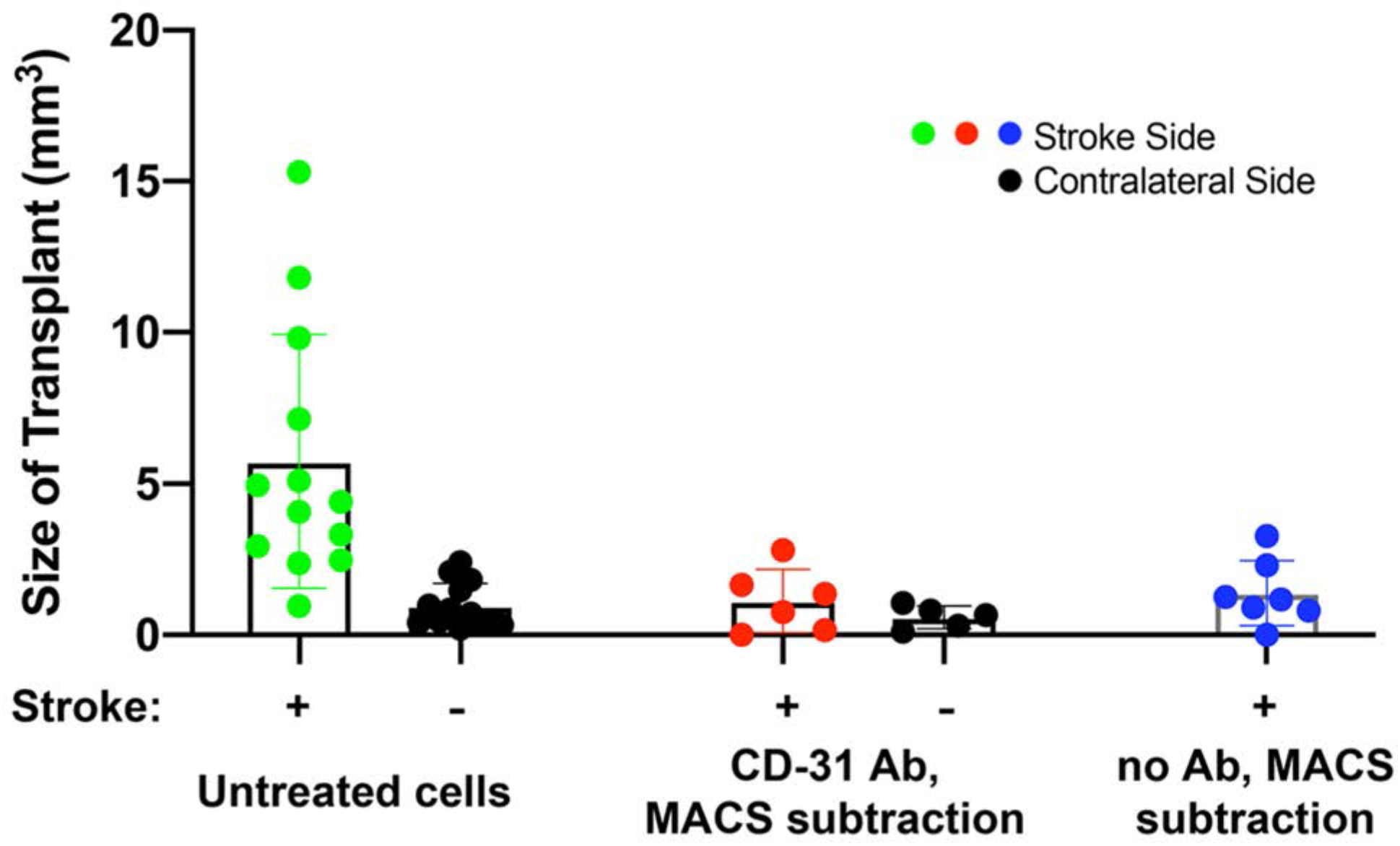
MACS procedure reduces the size of grafts regardless of the presence or not of vascular cells. Size of transplants without treatment (data taken from Figure 4C), with MACS immuno-exclusion of CD31+ cells, and with MACS preparation without antibody. N = 13 mice without treatment, N = 6 mice with CD31 MACS, N = 7 with no antibody with MACS.

**Supplementary Figure 4.**
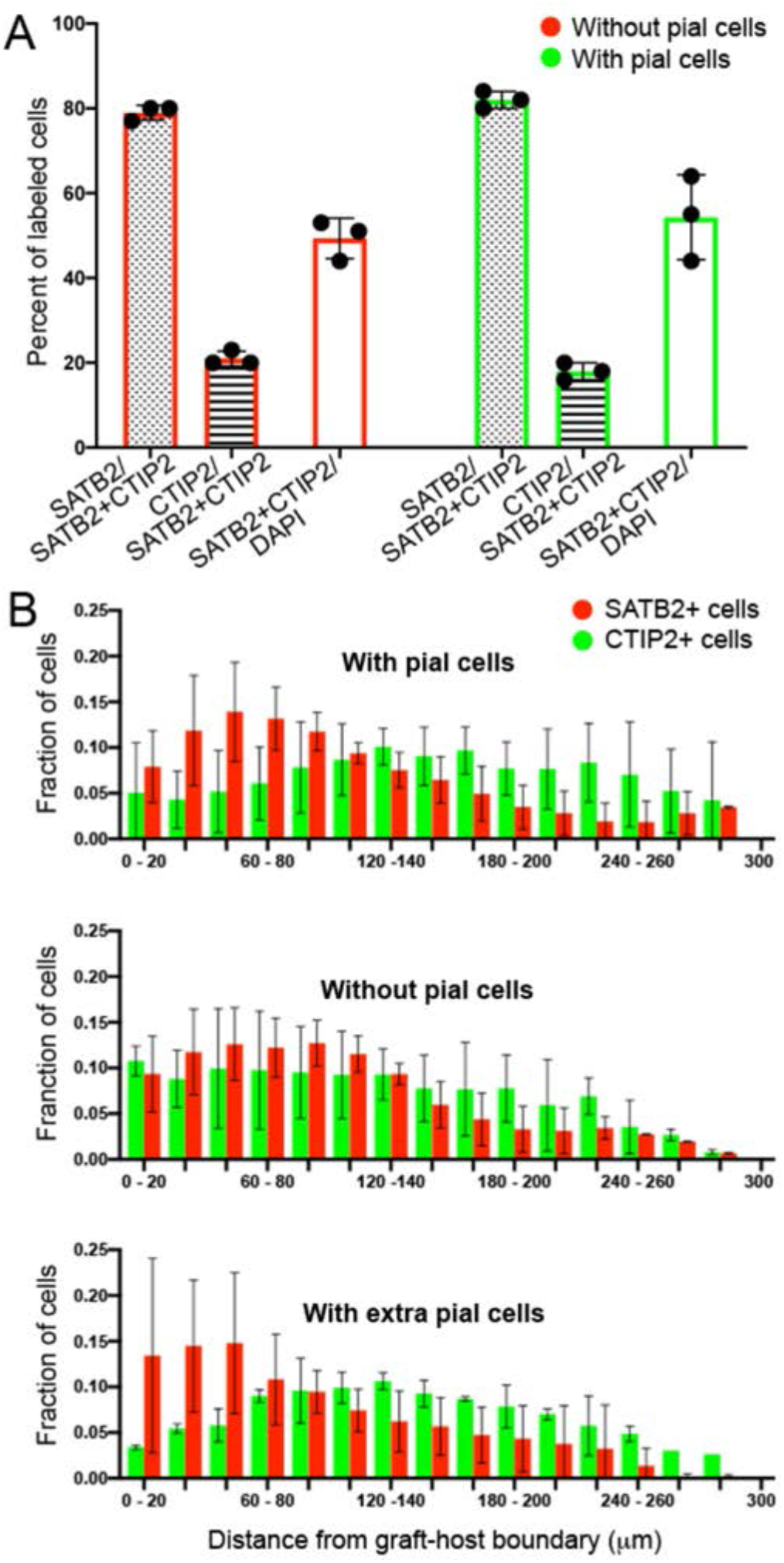
Exclusion of pial cells does not affect neuronal composition, but may reduce pseudo-layering. **A)** The ratios of SATB2+, CTIP2+, and total neurons is unaffected by the including pial cells in the grafts. **B)** Pseudo-layering appears to correlate with the presence of pial cells, despite being added as dissociated mixed cells. N = 4 mice with pial cells, 3 without pial cells, and 3 with extra (2-3X more) pial cells.

